# Adherens junctions limit the extent of the septate junctions in *Drosophila* midgut enterocytes but are not required for apical-basal polarity

**DOI:** 10.1101/2024.03.22.586295

**Authors:** Cátia A. Carvalho, Mihoko Tame, Daniel St Johnston

**Affiliations:** The Gurdon Institute and the Department of Genetics, University of Cambridge, Tennis Court Rd, Cambridge CB2 1QN; Department of Cell and Developmental Biology, School of Medicine, Faculty of Medical and Health Sciences, Tel Aviv University, Tel Aviv 69978, Israel

## Abstract

Adherens junctions formed by E-cadherin adhesion complexes play central roles in the organisation and apical-basal polarisation of both mammalian and insect epithelia. Here we investigate the function of the components of the E-cadherin adhesion complex in the *Drosophila* midgut epithelium, which establishes polarity by a different mechanism from other fly epithelia and has an inverted junctional arrangement, in which the adherens junctions lie below the septate junctions. Unlike other epithelial tissues, loss of E-cadherin, Armadillo (β-catenin) or α-catenin has no effect on the polarity or organisation of the adult midgut epithelium. This is not due to redundancy with N-cadherin, providing further evidence that the midgut polarises in distinct way from other epithelia. However, *E-cadherin* (*shg*) and *armadillo* mutants have an expanded septate junction domain and a smaller lateral domain below the septate junctions. Thus, E-cadherin adhesion complexes limit the basal extent of the septate junctions. This function does not appear to depend on the linkage of E-cadherin to the actin cytoskeleton because α-catenin mutants do not significantly perturb the relative sizes of the septate and sub-septate junction domains.

## Introduction

A defining feature of epithelial cells is their ability to adhere to each other laterally to form epithelial sheets and tubes. In most epithelia, this lateral adhesion depends on adherens junctions formed by the E-cadherin adhesion complex, composed of the homophilic adhesion protein, E-cadherin (encoded by the *shotgun* gene) and the cytoplasmic adaptor proteins, β-catenin (Armadillo in *Drosophila*), α-catenin and P120 catenin (Harris and Tepass, 2010; Roy and Berx, 2008; Tepass et al., 2001). E-cadherin binds in trans through opposing extracellular cadherin repeats (EC1-4) in neighbouring cells and also interacts with other E-cadherin molecules in cis to form clusters (Mège and Ishiyama, 2017). P120 catenin binds to the cytoplasmic, juxtamembrane domain of E-cadherin to increase adhesion and antagonise E-cadherin endocytosis, and this is essential for robust cell-cell adhesion in vertebrates but is dispensable in *Drosophila* (Davis et al., 2003; Ireton et al., 2002; Pacquelet and Rørth, 2005; Pacquelet et al., 2003; Thoreson et al., 2000). The E-cadherin cytoplasmic domain contains a more C-terminal binding site for β-catenin, which in turn recruits α-catenin, which links the complex to the F-actin cytoskeleton (Desai et al., 2013; Harris and Tepass, 2010).

The establishment of E-cadherin-dependent adherens junctions is intimately linked to the development of polarised epithelial cells in vertebrates. For example, expressing E-cadherin in unpolarised fibroblasts is sufficient to induce the polarised trafficking of basolateral proteins to cell contact regions (McNeill et al., 1990). Furthermore, loss of E-cadherin in mouse embryos leads to a failure to form the trophectoderm epithelium and arrests development, while the specific deletion of E-cadherin in the skin not only removes the adherens junctions but causes defects in the overlying tight junctions and a loss of barrier function (Larue et al., 1994; Roy and Berx, 2008; Tunggal et al., 2005). Similarly, siRNA knockdown of α-catenin in MDCK cells disrupted polarity completely (Capaldo and Macara, 2006).

The organisation of junctions in most invertebrate epithelia is inverted compared to that in vertebrates, as the adherens junctions concentrate in zonula adherens at the top of the lateral domain above more basal septate junctions, which are analogous to vertebrate tight junctions and perform the barrier function (St Johnston and Ahringer, 2010; Tepass et al., 2001). Nevertheless, loss of E-cadherin complex components has a similar effect on epithelial organisation and polarity as in vertebrates. It is not possible to remove all E-cadherin from the *Drosophila* embryo because there is a large maternal contribution that is required for normal oogenesis (Tepass et al., 2001). However, in embryos derived from germline clones of strong β-catenin mutants (*arm*) that also lack zygotic β-catenin, embryonic tissues lose their epithelial characteristics at gastrulation, producing unpolarised mesenchymal cells (Cox et al., 1996). A similar phenotype is observed with RNAi-mediated knock down of both maternal and zygotic α-catenin (Sheppard et al., 2022). The role of E-cadherin adhesion complexes has also been examined in the follicular epithelium that surrounds developing female germline cysts (Tepass et al., 2001). E-cadherin mutant clones cause various developmental defects, but do not disrupt epithelial organisation or polarity because of redundancy with N-cadherin (Godt and Tepass, 1998; González-Reyes and St Johnston, 1998; Pacquelet and Rørth, 2005; Tanentzapf et al., 2000). By contrast, *armadillo* and α*-catenin* mutant clones lose their epithelial organisation, flatten and have disrupted apical-basal polarity (Bonello et al., 2021; Sarpal et al., 2012; Tanentzapf et al., 2000).

E-cadherin adhesion complexes contribute to epithelial polarity by several mechanisms. First, by linking sites of cell adhesion to the actin cytoskeleton, they generate robust intercellular junctions that can resist the forces exerted during morphogenesis (Harris and Tepass, 2010b; Lecuit and Yap, 2015). Second, the key polarity factor Bazooka (Par-3) is recruited to adherens junctions through direct interactions with E-cadherin and β-catenin, where it is thought to form phase separated condensates that provide a barrier between the apical and lateral domains (Bonello et al., 2021; Buckley and St Johnston, 2022; Kono et al., 2019; Liu et al., 2020; Wei et al., 2005). Third, RhoGAP19D is recruited to adherens junctions along the lateral domain, where it inactivates Cdc42, thereby restricting the activity of the apical Cdc42/Par-6/aPKC complex to the apical side of the cell (Fic et al., 2020).

The intestinal epithelium of the *Drosophila* midgut has a different organisation from other fly epithelia as it forms septate junctions at the top of the lateral domain above lateral adherens junctions, a junctional arrangement that resembles that in vertebrates (Baumann, 2001). Furthermore, the midgut enterocytes do not express key polarity factors, such as Bazooka (Par-3), and their apical-basal polarisation does not require any of the canonical epithelial polarity factors that polarise other *Drosophila* epithelia (Chen et al., 2018). These differences may reflect the endodermal origin of the midgut, or the fact that, unlike other fly epithelia, enterocytes polarise in a basal to apical direction as they differentiate from basally-located enteroblasts and integrate into the epithelium (Chen and St Johnston, 2022; Galenza et al., 2023). The distinct organisation of the midgut epithelium raises the question of what roles E-cadherin adhesion complexes play in enterocyte polarisation. E-cadherin has previously been shown to mediate adhesion between basal intestinal stem cells (ISCs) and their progeny enteroblasts (EBs), where it acts to delay EB detachment to allow Delta-Notch signalling to specify EB identity (Maeda et al., 2008). In addition, down-regulation of E-cadherin in apoptotic enterocytes triggers the release of epidermal growth factors to stimulate compensatory divisions of the ISCs, thereby maintaining homeostasis (Liang et al., 2017). E-cadherin must also be removed from the apical surface of integrating enteroblasts to establish an apical membrane initiation site (AMIS) (Chen and St Johnston, 2022). However, the effects of E-cadherin complex depletion on enterocyte polarity and differentiation have not been analysed. Here we set out to investigate the role of E-cadherin complex components in this process and demonstrate that adherens junctions are dispensable for enterocyte polarity but play a role in defining the size and position of the septate junctions and the extent of the lateral domain.

## Results

### Adherens junction components localize below the septate junctions

To verify the composition of adherens junctions in the *Drosophila* midgut, we used antibodies and protein trap lines to examine the localization of the E-cadherin adhesion complexes components (Fig. 1A). Endogenously-tagged E-cadherin-GFP is highly expressed in ISCs and EBs, as previously reported (Chen and St Johnston, 2022; Huang et al., 2009; Maeda et al., 2008). E-cadherin is also expressed in enterocytes, albeit at lower levels, and localizes strongly along the region of the lateral membrane below the septate junctions, which are marked by Coracle (Fig. 1B). In cross-sections through the epithelium, E-cadherin is distributed in diffraction-limited puncta along the cell membranes, which presumably correspond to spot adherens junctions. E-cadherin is not restricted to the lower portion of the lateral domain, however, as low levels are also found throughout the septate junction region (Fig. 1C). Antibody staining for β**-**catenin (Armadillo) and an α-catenin protein trap line displayed similar distributions to E-cadherin (Fig. 1D,E). By contrast, P120-catenin is not expressed at detectable levels in the midgut, since a monoclonal antibody that recognizes P120ctn in other tissues gave only non-specific staining of the septate junctions, which was still present in *p120ctn* hmozygous clones (Iyer et al., 2019; Magie et al., 2002). These results confirm that Adherens junctions form below the septate junctions in the midgut, which is the opposite way round to other *Drosophila* epithelia but resembles the junctional arrangement in vertebrates (Chen et al., 2018).

**Figure 1.**
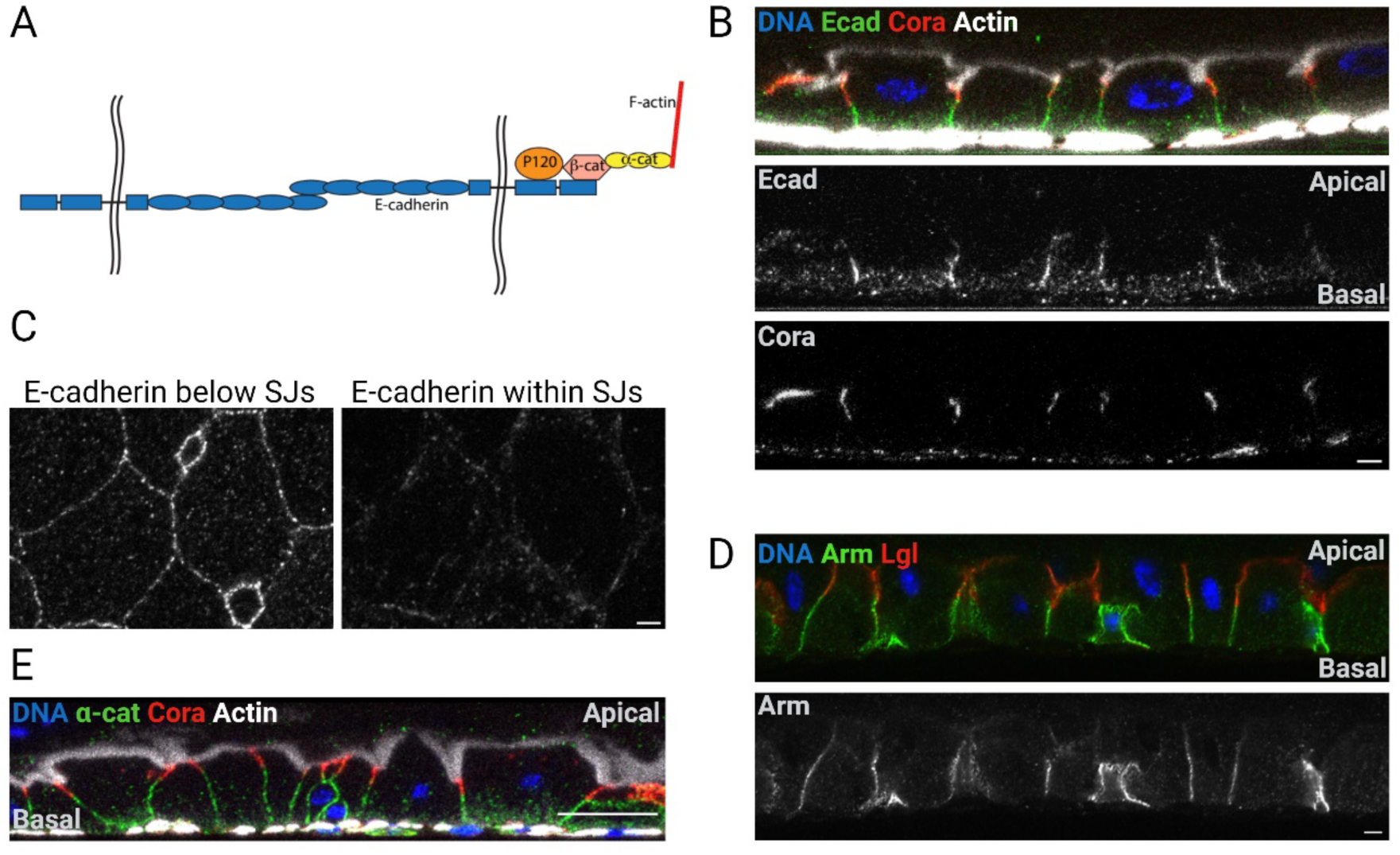
E-cadherin, β-catenin and α-catenin co-localize beneath the Septate junctions in the *Drosophila* midgut. **A.** A diagram showing the components of E-cadherin adhesion complexes. **B.** The localisation of endogenously tagged E-cadherin-GFP (green) along the apical-basal axis of the midgut epithelium, stained for DNA (DAPI; blue), Coracle (red) and F-actin (white). **C.** A horizontal view of E-cadherin at the level of the Adherens junctions (left) and at the level of the septate junctions (right). **D.** β-catenin localization (green) along the apical basal axis of the midgut epithelium, stained for DNA (DAPI; blue) and Lgl (red). **E.** An apical-basal confocal section showing the localisation of an α-catenin protein trap line (green) stained for DNA (DAPI; blue), Coracle (red) and actin (white). The scale bars in B-E = 20 µm.

### The localisations of Adherens junctions components are mutually-dependent

The core components of the adherens junctions rely on mutually-dependent interactions to localize at the plasma membrane. Although intercellular adhesion is nucleated by the trans homophilic interaction between E-cadherin extracellular domains, E-cadherin’s efficient trafficking from the endoplasmic reticulum to the plasma membrane depends on the binding of its cytoplasmic tail to β-catenin (Chen et al., 1999; Curtis et al., 2008; Tanentzapf et al., 2000). Conversely, loss of E-cadherin impairs the targeting of β-catenin and α-catenin to the plasma membrane, and α-catenin depletion decreases both E-cadherin and β-catenin localization at cell-cell contact sites (Bonello et al., 2021; Capaldo and Macara, 2006; Pacquelet and Rørth, 2005; Sarpal et al., 2012). To investigate if these mutual dependencies are maintained in the *Drosophila* midgut, we generated homozygous clones for null alleles in β-catenin (*arm^YD35^*) E-cadherin (*shg^R69^*) and α-catenin (*α-cat^1^*) using the MARCM system (Godt and Tepass, 1998; Lee and Luo, 2001; Peifer and Wieschaus, 1990; Sarpal et al., 2012). As expected, *arm^YD35^* homozygous enterocytes lack β-catenin staining at the membrane, which is outlined by staining for the septate junction protein, Mesh, (Fig. 2A). A plot of the β-catenin signal intensity across the lateral membrane in the *arm^YD35^*/+ heterozygous control shows a clear peak at the cell junction, whereas the homozygous mutant clone shows a uniform level of staining. Since *arm^YD35^*is a null allele that produces no protein(Peifer and Wieschaus, 1990), this residual signal may correspond in part to perdurance of the wild-type protein after clone induction, but is most likely background staining (Fig. 2B,G). GFP-labelled *shg^R69^* homozygous enterocytes also lose β-catenin staining from the plasma membrane (Fig. 2C). β-catenin intensity plots show no peak at the adherens junction, indicating that E-cadherin is required for all β-catenin localization at the membrane (Fig. 2D & G). Furthermore, the cytoplasmic signal remains at similar levels to that in *shg^R69^*/+ heterozygous cells, suggesting that the β-catenin that fails to localise to junctions is degraded, most probably by the APC/Axin destruction complex (Fig. 2H) (Clevers, 2006). Similar results were observed with additional *shotgun* and *armadillo* alleles (Fig. S1 A-E). Lastly, mutant enterocytes lacking α-catenin displayed reduced levels of β-catenin at the lateral membrane, but still had a clear enrichment at the junctions (Fig. 2E,G). β-catenin intensity plots across mutant enterocytes reveal a more uniform distribution of β-catenin throughout the cell, with slightly higher cytoplasmic levels than in the heterozygous controls (Fig. 2F,H). Thus, the mutual dependencies between the components of the E-cadherin adhesion complex are similar to those in other tissues, despite the altered junctional arrangement in the midgut.

**Figure 2:**
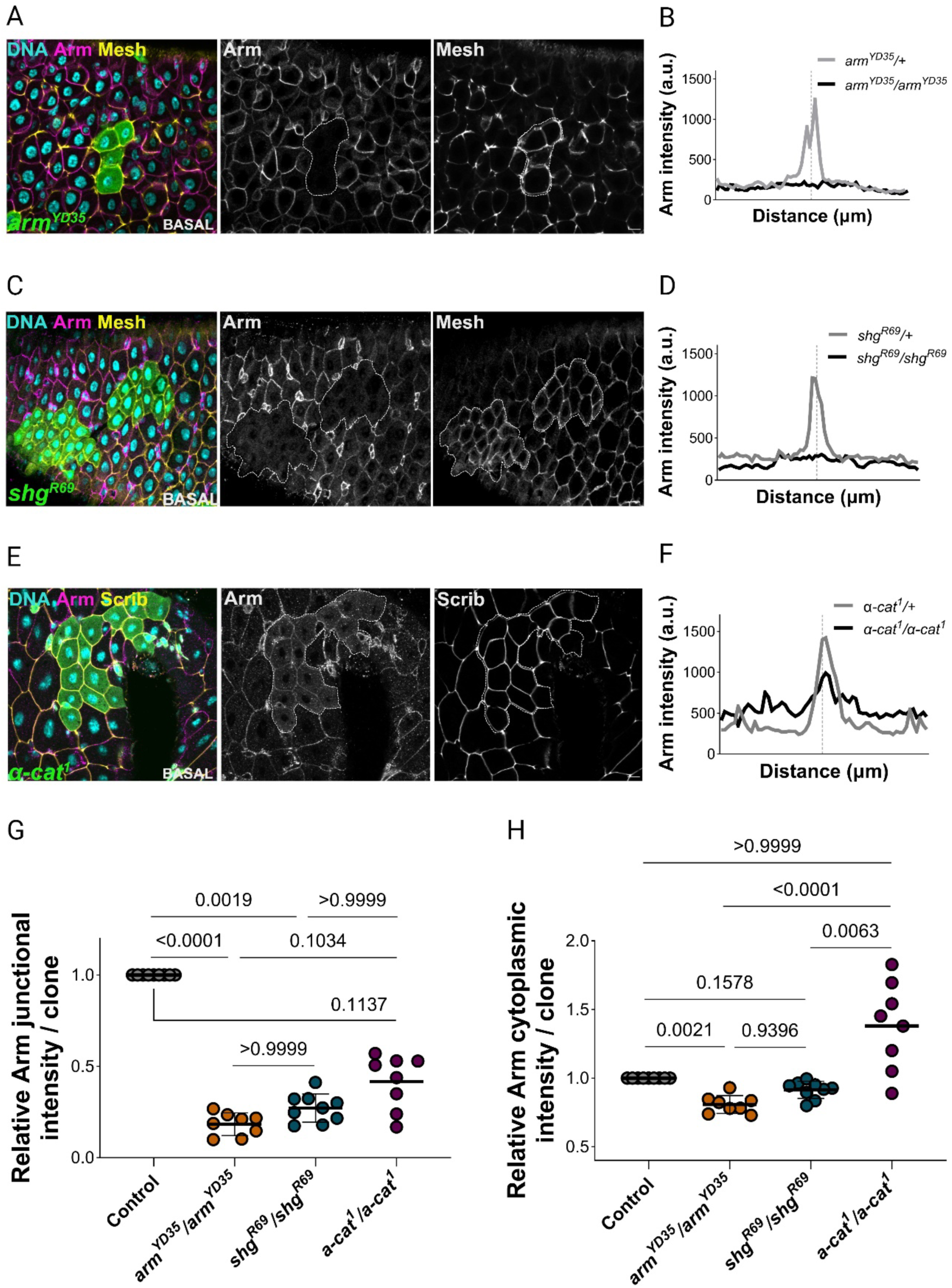
The Adherens junctions core components are mutually dependent for their membrane localization. **A**. A horizontal view of a clone of *arm^YD35^* homozygous cells (GFP-positive; green) stained for DNA (DAPI; blue), Armadillo (magenta) and Mesh (yellow). The clone boundaries are outlined by white dotted lines; Scale bar=10 µm. **B.** Plots of Armadillo signal intensity across two cells, centered at the position of the cell-cell boundary (dotted vertical line). The black line shows data from *arm^YD35^*homozygous enterocytes and the grey line, data from *arm^YD35^*/+ heterozygous enterocytes; **C**. A horizontal view of a representative *shg^R69^*homozygous clone marked by GFP expression (green), stained for DNA (DAPI; blue), Armadillo (magenta) and Mesh (yellow); clone boundaries are outlined in white. **D.** Armadillo intensity plots in *shg^R69^* homozygous enterocytes (black line) and *shg^R69^/+* heterozygous enterocytes (grey line). **E.** A horizontal view of a *α-cat^1^* mutant clone (GFP) stained for DNA (DAPI; blue), Armadillo (magenta) and Scribble (yellow. **F.** Armadillo intensity plots in *α-cat^1^* homozygous (black line) and heterozygous (grey line) cells. **G.** Measurements of Armadillo junctional intensity at the AJs in heterozygous control cells (standardized mean=1, 115 cell-cell junctions), *arm^YD35^/arm^YD35^* (mean=0.18±0.06, 91 cell-cell junctions), *shg^R69^/shg^R69^* cells (mean= 0.27±0.08, 140 cell-cell junctions) and *α-cat^1^/α-cat^1^* cells (mean=0.42±0.15, 103 cell-cell junctions). **H.** Measurements of Armadillo intensity in the cytoplasm in heterozygous controls (standardized mean=1, 93 cells), *arm^YD35^/arm^YD35^* (mean=0.81±0.07, 70 cells), *shg^R69^/shg^R69^* (mean=0.91±0.06, 119 cells) and *α-cat^1^/α-cat^1^* (mean=1.38±0.32, 92 cells) enterocytes. The scale bars in A, C and E = 10µm.

### Adherens junctions are not required for the establishment of apical-basal polarity

Unlike other *Drosophila* epithelia, the apical-basal axis of *Drosophila* midgut is established in a basal-to-apical direction as enteroblasts integrate into the epithelium, and the adherens junctions therefore form first before the more apical septate junctions (Chen and St Johnston, 2022; Chen et al., 2018). Given the key role of adherens junctions in the apical-basal polarity of other epithelia in both vertebrates and *Drosophila*, we investigated whether loss of the E-cadherin adhesion complex would disrupt enteroblast/enterocyte integration or polarisation. We generated mutant stem cell clones lacking each adherens junction component and examined their phenotypes 3-15 days after clone induction to reduce the perdurance of pre-existing wild-type protein and to ensure that the cells lacked the protein of interest from birth. Loss of E-cadherin did not prevent mutant enteroblasts from integrating into the epithelium and differentiating as enterocytes (Fig. 3A-A’). Longitudinal views show that *shg^R69^/shg^R69^* enterocytes have well-defined apical domains contacting the gut lumen, localise Canoe normally at the apical-lateral boundary and are properly attached to the basement membrane (Fig. 3A’’). Like *shg^R69^/shg^R69^*cells, *arm^YD35^/arm^YD35^* mutant cells integrate into the epithelium and form a normal apical brush border, marked by the localisation of atypical protein kinase C (aPKC), and septate junctions labelled by Coracle (Fig. 3 B-B’’). *α-catenin* null cells also integrate and establish a wild-type apical-basal axis, with a fully-formed basal labyrinth, marked by Nervana (Nrv)(Fig. 3 C-C’’). To verify these results, we knocked-down the core components of the E-cadherin complex by RNAi in enterocytes throughout the entire midgut. Downregulation of E-cadherin using two different RNAi lines led to a marked reduction in β-catenin localisation to cell junctions but did not cause any apical-basal polarity defects (Fig. S2 A-C). Downregulation of β-catenin or α-catenin also had no effect on tissue organisastion or polarity (Fig. S2 A,D-E). Altogether, these results demonstrate that the core components of the adherens junctions are not required for either the establishment or maintenance of apical-basal polarity in the midgut epithelium, unlike all other epithelia that have been examined so far.

**Figure 3:**
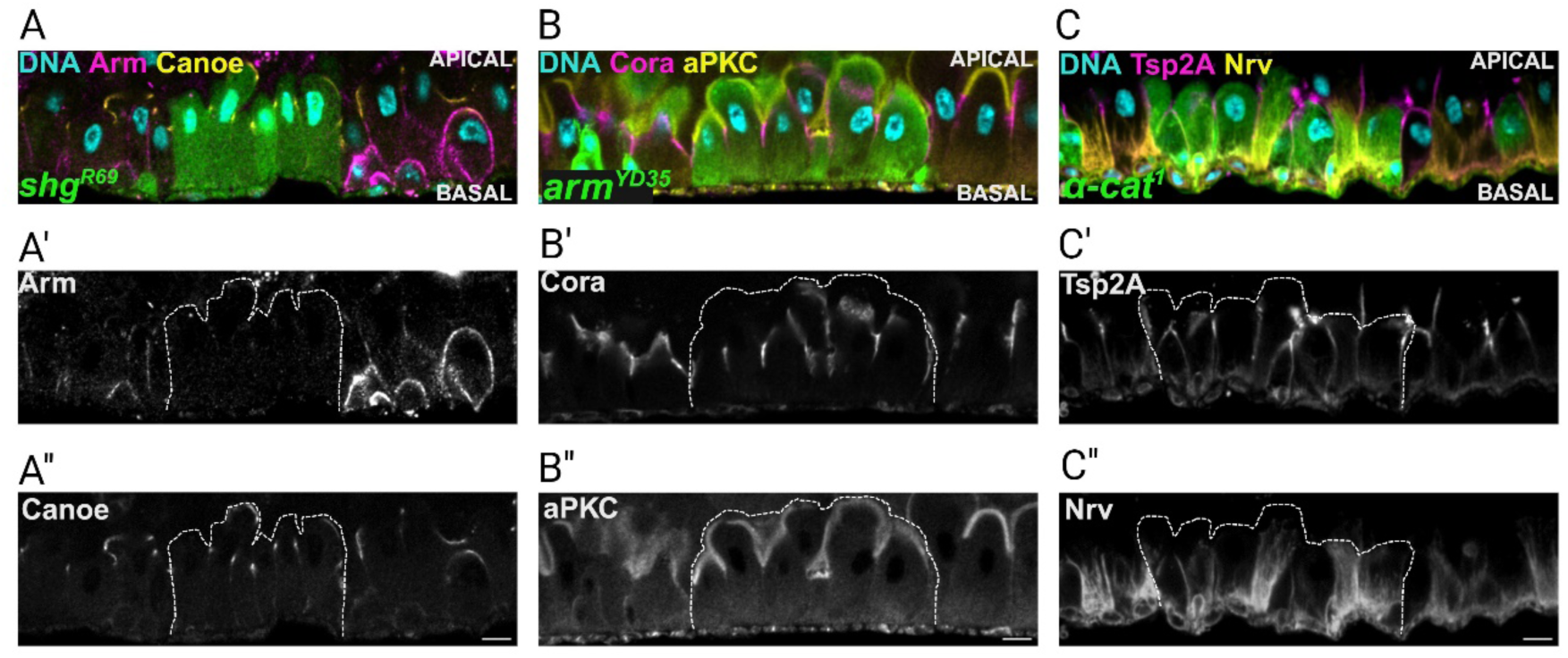
Adherens junctions are not required for apical-basal polarity in the midgut. **A.** An apical-basal section showing a *shg^R69^* homozygous clone (GFP; green) stained for DNA (DAPI; blue) (**A**), Armadillo (magenta and **A’**) and Canoe (yellow and **A’’**); the clone is outlined by a white dotted lines in **A’** and **A’’**;; **B.** An apical-basal section showing an *arm^YD35^* homozygous clone (GFP; green), stained for Coracle (magenta and **B’**) and aPKC (yellow and **B’’**); white dashed lines outline the clone in **B’** and **B’’**. **C.** An *α-cat^1^* homozygous clone (GFP; green) stained for Tsp2A (magenta and **C’**) and Nervana (yellow and **C’’**). The scale bars = 10 µm.

### E-cadherin and β-catenin regulate enterocyte shape independently of α-catenin

Although the adherens junctions are not required for enterocyte polarity, they could still play a role in regulating cell shape and cytoskeletal organisation, as they do in other epithelia (Bonello et al., 2021). To explore how E-cadherin-based adhesions modulate enterocyte morphology, we measured the circularity of mutant cells at the basal side as a proxy for the degree to which they adopted the normal polygonal shape of midgut epithelial cells. Enterocytes mutant for E-cadherin (0.77±0.11) became significantly more rounded than their heterozygous neighbours (0.70±0.12) (Fig. S3A,D). Similarly, loss of β-catenin (0.77±0.14) also significantly reduced the polygonal shape of the enterocytes compared to the controls (0.70±0.11) (Fig. S3B,D). Enterocytes mutant for α-catenin (0.76±0.13) were also rounder than the neighbouring enterocytes (0.71±0.12), but this difference was not statistically significant (Fig. S3C,D). Altogether, these data indicate that the loss or reduction in adhesion in the basal portion of the lateral domain allows mutant cells to become more circular.

### E-cadherin and β-catenin limit the basal extent of the Septate junctions

We next examined the contribution of the adherens junctions to lateral domain identity. The lateral domain of the enterocytes comprises two sub-domains: an apical septate junction domain and a more basal domain that normally contains adherens junctions, which we will refer to as the sub-septate junction domain. The septate junction domain is significantly extended in enterocytes depleted of E-cadherin (*shg^R69^)* (11.9±3.6µm) compared to the neighbouring heterozygous cells (7.8±2.7µm) (Fig. 4A,D). Similarly, enterocytes without β-catenin have longer septate junctions (10.7±4.6µm) relative to control cells (6.6±1.9µm) (Fig. 4 B,D). Cells lacking α-catenin also have slightly longer septate junctions (7.1±3.0µm versus 6.1±2.7µm), but this difference was not significant (Fig. 4 C,D). Thus, lateral E-cadherin adhesion complexes limit the basal extent of the septate junctions. Consistent with this, the domain of the lateral membrane below the septate junctions (the sub-SJ domain) is significantly shorter in both *shg^R69^/shg^R69^* enterocytes (9.6±3.6µm versus 18.2±4.1µm) and in *arm^YD35^/arm^YD35^*enterocytes (12.4±5.1µm versus 17.9±5.6µm) (Fig. 4 A,B,E). This decrease in the sub-septate junction domain is not observed upon loss of function of α-catenin (15.0±3.7µm versus 15.2±5.5µm) (Fig. 4 C,E). The strong reduction in the sub-septate junction domain in *shg^R69^/shg^R69^* enterocytes more than compensates for the longer septate junctions in these cells, leading to a significant decrease in the overall length of the lateral domain (21.5±5.4µm) compared to neighbouring heterozygous cells (26.0±4.6µm) (Fig. 4A,F). The reduction in total lateral domain length was not significant in enterocytes that lacked β-catenin or α-catenin, consistent with their weaker effects on the size of the sub-septate junction domain (Fig. 4B,C,F). To confirm that the reduction in the length of the sub-septate junction domain and the concomitant basal expansion of the septate junctions was caused by the mutations in *shotgun* and *armadillo*, and not by second hits on the mutant chromosomes, we repeated these experiments with another null allele of each gene (*shg^IG29^* and *arm^XP33^*). Consistent with our previous results, both *shg^IG29^* and *arm^XP33^*mutant cells showed a significant increase in the extent of the septate junction domain and a decrease in the length of the lateral membrane below the septate junctions (Fig S4 A-D). However, only *shg^IG29^* mutant cells showed a significant reduction in total cell height, indicating that the loss of E-cadherin has a stronger effect on the lateral domain organisation than loss of either catenin.

**Figure 4:**
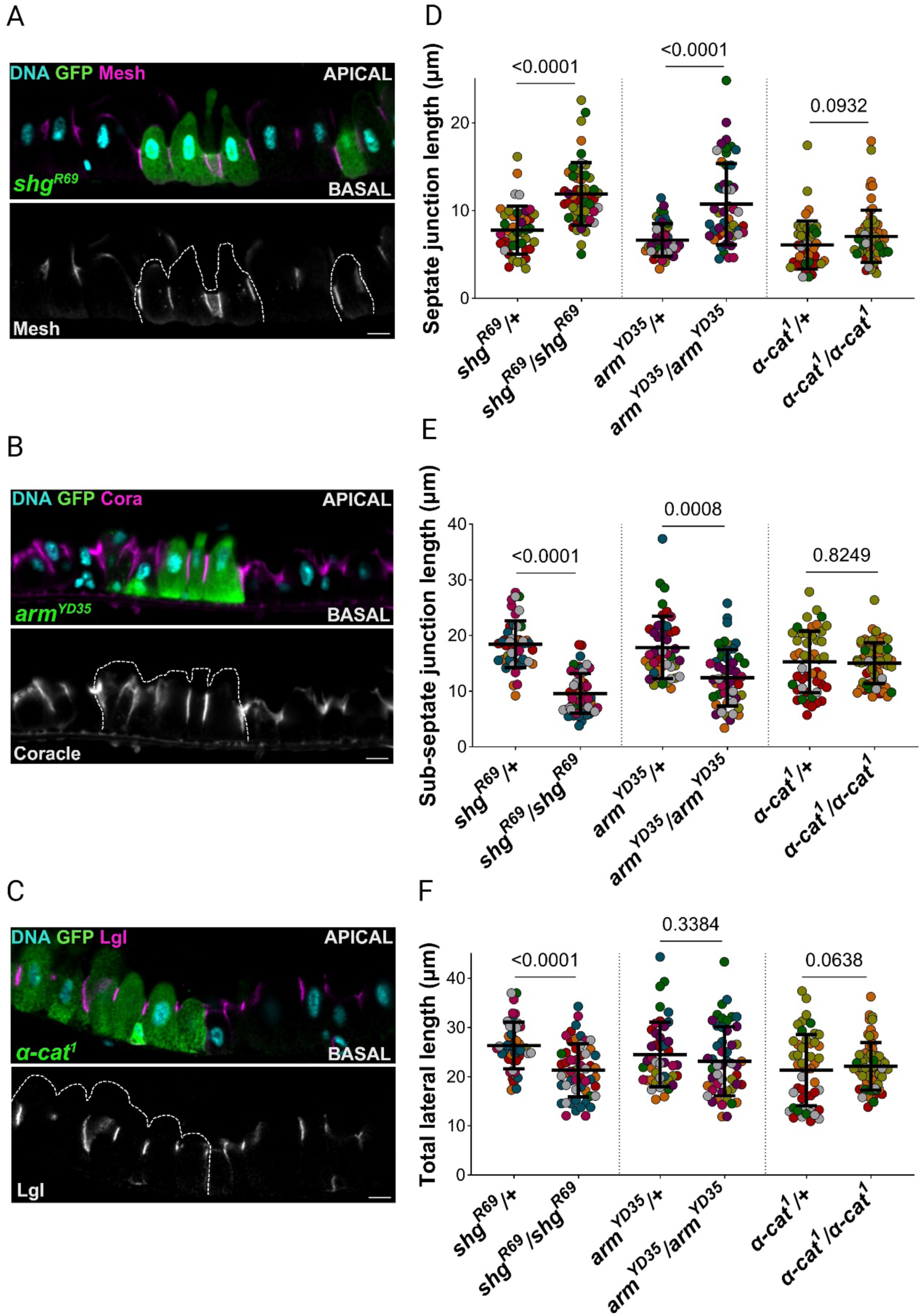
E-cadherin and β-catenin limit the extent of the Septate junctions. **A.** An apical-basal section through a midgut containing a *shg^R69^* homozygous clone marked by GFP expression (green) stained for DNA (DAPI, blue) and Mesh (magenta); the mutant cells are outlined in white in the lower panel. **B.** An *arm^YD35^* homozygous clone (GFP; green) stained for DNA (DAPI; blue) and Coracle (magenta). **C.** An *α-cat^1^* homozygous clone (GFP; green) stained for DNA (DAPI; blue) and Lgl (magenta); **D.** Measurements of septate junction length in *shg^R69^/*+ (39 junctions), *shg^R69^/shg^R69^* (46 junctions), *arm^YD35^/+* (44 junctions), *arm^YD35^/arm^YD35^* (50 junctions), *α-cat^1^/+* (47 junctions) and *α-cat^1^/α-cat^1^* (61 junctions) enterocytes. **E.** Measurements of the length of the sub-septate junction domain in the same cells as in **D.**; **F.** Measurements of total lateral length of the same cells as in **D.** The scale bars = 10 µm.

### Loss of N-cadherin stabilises β-catenin in enterocytes

One possible reason for the lack of a clear epithelial polarity phenotype in E-cadherin null cells is redundancy with N-cadherin, as is the case in the follicular epithelium that surrounds developing *Drosophila* egg chambers (Straub et al., 2011; Tanentzapf et al., 2000). To test this possibility, we generated clones of cells homozygous for a strong allele of N-cadherin. Mutant cells integrate normally into the midgut epithelium and differentiate into fully-polarised enterocytes with normal polygonal shapes, as shown by their circularity (0.67±0.14 versus 0.72±0.11) (Fig. 5A,B and Fig. S4E). We confirmed these results by depleting N-cadherin by RNAi in the enterocytes throughout the entire midgut and observed wild-type epithelial polarity and organisation (Fig S2A,F). Furthermore, the loss of N-cadherin had no effect on the lengths of the septate junctions (8.4±2.7µm versus 7.1±2.8µm), the sub-septate junction domain (20.2±6.2µm versus 19.3±4.0µm) or the entire lateral domain (28.6±7.0µm versus 26.4±4.7µm) (Fig. 5B,C). Unlike E-cadherin mutants, however, enterocytes depleted for N-cadherin showed significantly increased levels of β-catenin at the adherens junctions, relative to the heterozygous control cells (Fig. 5 B,D). *Ncad^405^/Ncad^405^*enterocytes also have increased levels of β-catenin in the cytoplasm (Fig. 5 A,D). These results demonstrate that loss of N-cadherin somehow stabilizes β-catenin, possibly because E-cadherin is up-regulated to compensate for the loss of N-cadherin and stabilises β-catenin.

**Figure 5:**
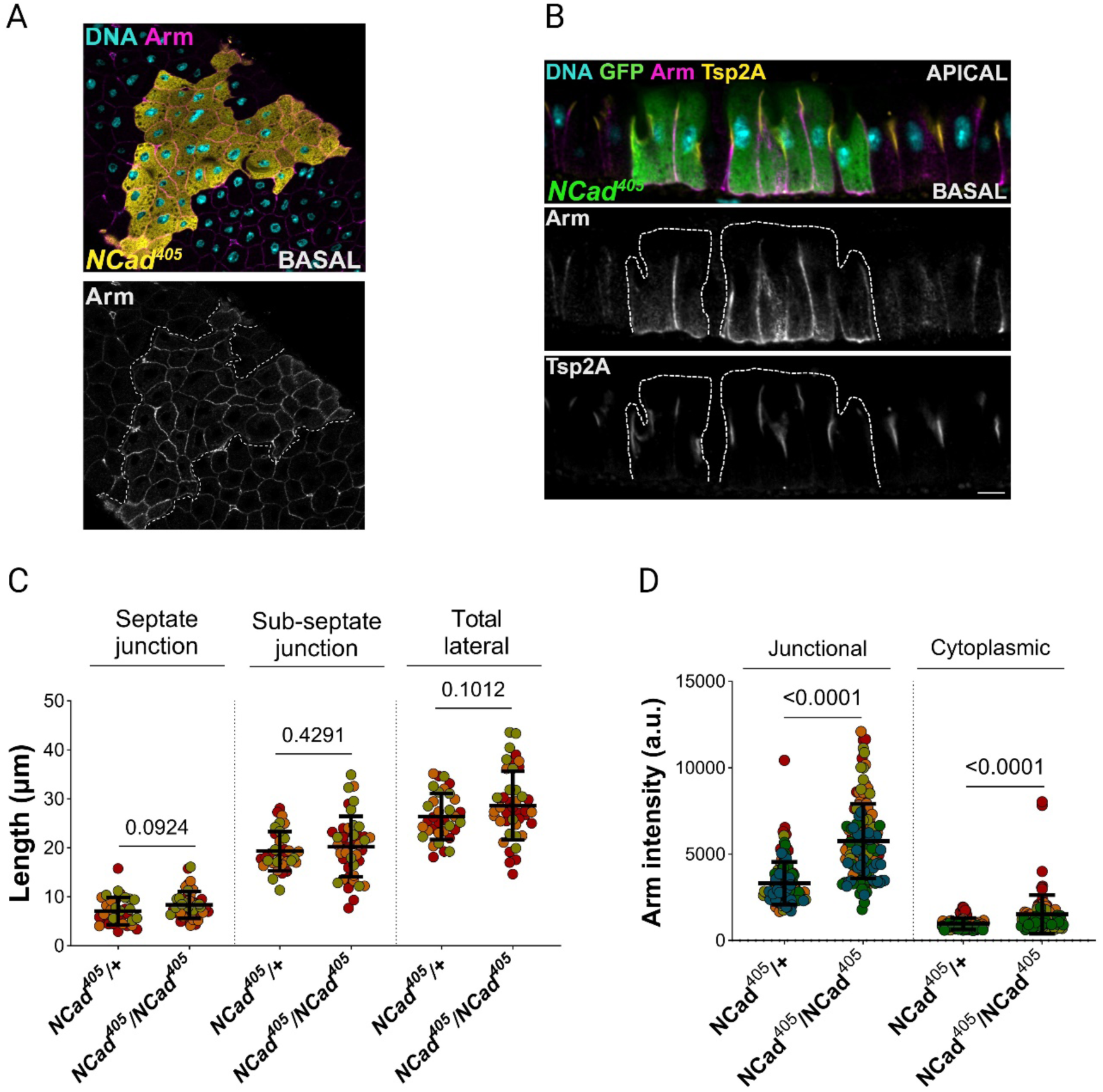
Loss of N-cadherin stabilizes β-catenin but does affect enterocyte polarity or shape. **A.** A horizontal section through a midgut containing a *Ncad^405^* homozygous mutant clone (yellow) stained for DNA (DAPI; blue) and Armadillo (magenta); the clone is highlighted in white in the lower panel. Scale bar=10 µm; **B.** An apical-basal section containing *Ncad^405^* (GFP) clones stained for DNA (DAPI; blue), Armadillo (magenta) and Tsp2A (yellow). **C.** Measurements of the lengths of the septate junctions, sub-septate junction domain and total lateral length in *Ncad^405^/+* (34 junctions) and *Ncad^405^/Ncad^405^* (43 junctions) enterocytes. **D.** Measurements of Armadillo intensity in the sub-septate junction domain (*Ncad^405^/+*: 3317±1233, 133 cell-cell junctions; *Ncad^405^/Ncad^405^*: 5750±2158, 136 cell-cell junctions) and in the cytoplasm (*Ncad^405^/+*: 975±332, 101 cells; *Ncad^405^/Ncad^405^*: 1516±1117, 93 cells).

### N-cadherin is not redundant with E-cadherin

If N-cadherin can compensate for the loss of E-cadherin in the enterocytes, the concomitant depletion of both cadherins should produce a stronger phenotype than removing either alone. We produced cells lacking both cadherins by inducing MARCM clones homozygous for a null allele of E-cadherin (*shg^IG29^*), in which N-cadherin was also knocked down by RNAi. These GFP-positive cells lacking both cadherins from birth still integrated into the epithelium and displayed normal apical-basal polarity (Fig. 6A,B and Fig. S2A,F). Thus, N-cadherin does not compensate for the loss of E-cadherin and both cadherins are dispensable for epithelial polarity and organisation in the midgut. Indeed, depleting N-cadherin from E-cadherin mutant clones partially suppresses the extension of the septate junctions observed in *shg^IG29^* homozygous clones but does not rescue the reduction of the length of the entire lateral domain induced by E-cadherin loss (Fig. 6B,C). Upon loss of E- and N-cadherin, β-catenin no longer localised to the sub-septate junction domain, as expected (Fig. 6B,D). Nevertheless, the levels of β-catenin were increased in the cytoplasm of E- and N-cadherin deficient cells compared to control cells (Fig. 6A,B,D). This phenotype is also seen when only N-cadherin is mutant (*NCad^405^*), ruling out the possibility that the higher levels of non-junctional β-catenin in the latter are due to the compensatory up-regulation of E-cadherin. Thus, N-cadherin plays a role in the destabilisation of β-catenin that is independent of E-cadherin.

**Figure 6:**
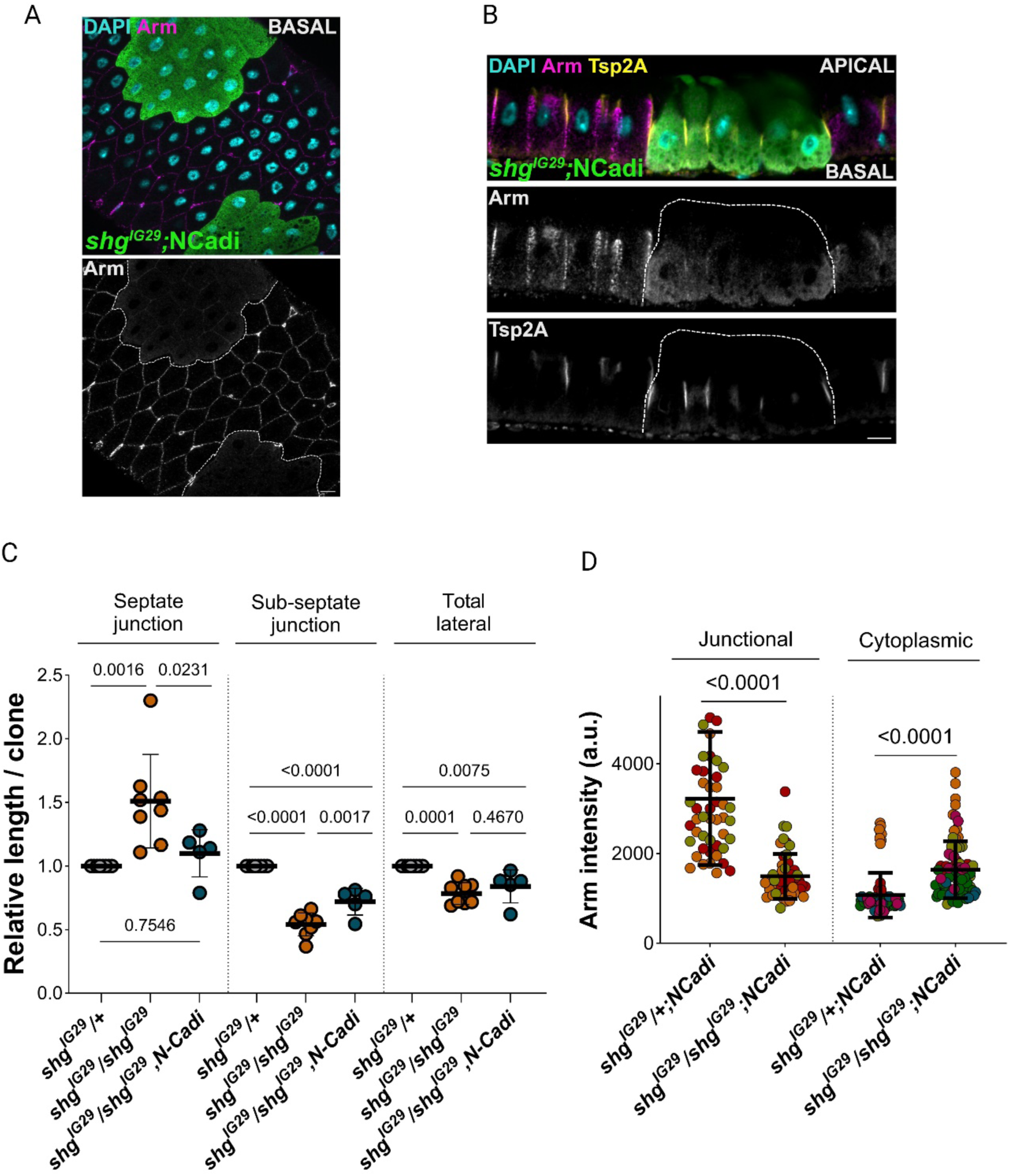
N-cadherin does not compensate for loss of E-cadherin in the midgut epithelium. **A.** A horizontal section though a midgut containing *shg*^IG29^ mutant clones expressing Ncad RNAi (GFP; green), stained for DNA (DAPI; blue) and Armadillo (magenta); the clones are outlined in white in the lower panel; Scale bar = 10 µm; **B.** An apical-basal section through a midgut containing a *shg*^IG29^ mutant clone expressing Ncad RNAi (GFP; green), stained for DNA (DAPI; blue), Armadillo (magenta) and Tsp2A (yellow). Scale bars = 10 µm; **C.** Measurements of the lengths of the septate junctions (*shg^IG29^/+* standardized mean=1, *shg^IG29^/shg^IG29^*mean=1.51±0.37, *shg^IG29^/shg^IG29^*, *Ncad RNAi* mean=1.10±0.19) the sub-septate junction domain (*shg^IG29^/+* standardized mean=1, *shg^IG29^/shg^IG29^* mean=0.54±0.09, *shg^IG29^/shg^IG29^*, *Ncad RNAi* mean=0.72±0.11) and the total lateral length (*shg^IG29^/+* standardized mean=1, *shg^IG29^/shg^IG29^*mean=0.78±0.09, *shg^IG29^/shg^IG29^*, *Ncad RNAi* mean=0.84±0.13) (*shg^IG29^/+ and shg^IG29^/shg^IG29^* as in Fig. S4B; *shg^IG29^/+, Ncad RNAi*: 47 junctions; *shg^IG29^/shg^IG29^, Ncad RNAi*: 53 junctions) in clones depleted for both cadherins and respective controls; **D.** Measurements of Armadillo intensity at the junctions (sub-septate junction domain; *shg^IG29^/+, Ncad RNA RNAi*: 48 junctions; *shg^IG29^/shg^IG29^, Ncad RNAi*: 52 junctions) and in the cytoplasm (*shg^IG29^/+,Ncad RNAi*: 85 cells; *shg^IG29^/shg^IG29^, Ncad RNAi*: 79 cells) of *shg^IG29^* heterozygous and homozygous enterocytes depleted for N-cadherin.

## Discussion

Our results demonstrate that adherens junctions are not required for the organisation or polarisation of the midgut epithelium. This contrasts with observations in other epithelia, where depletion of E-cadherin adhesion complexes disrupts epithelial organisation to produce mesenchymal-like cells and impairs apical-basal polarity (Bonello et al., 2021; Capaldo and Macara, 2006; Cox et al., 1996; Larue et al., 1994; Roy and Berx, 2008; Sheppard et al., 2022; Tunggal et al., 2005). This reinforces the evidence that the midgut epithelium polarises by a different mechanism from other *Drosophila* epithelia. Indeed, while ectodermal epithelia can polarise in the absence of septate junctions, which form late in epithelial differentiation, the septate junctions are essential for apical domain formation in the midgut (Chen et al., 2018; Laprise et al., 2009; Tepass et al., 2001). The septate junctions in the midgut form in the position that the adherens junctions occupy in other fly epithelia, namely at the boundary between the apical and lateral domains. Thus, the septate junctions seem to have taken over the role of the adherens junctions in apical-basal polarity in the midgut.

Although E-cadherin adhesion complexes are dispensable for polarity in the midgut, they do play a role in determining the relative sizes of the septate junction and sub-septate junction domains. Not much is known how the sizes of membrane domains are controlled, but this presumably (1) depends on the rates at which exocytosis delivers membrane and membrane proteins to the domain, (2) the rate at which endocytosis removes them and (3) the factors that define the boundaries between domains. The reduction in the size of the sub-septate junction domain is likely to be due at least in part to the failure to deliver exocytic vesicles containing E-cadherin. It is also possible that E-cadherin promotes the delivery of other proteins to this domain either by co-transport or by enhancing the delivery of other exocytic vesicles as observed in mammalian cells (Grindstaff et al., 1998; Yeaman et al., 2004). The exocyst associates with β-catenin and localises to adherens junctions in the *Drosophila* pupal notum (Langevin et al., 2005). Thus, E-cadherin/β-catenin complexes could enhance secretion to the sub-septate junction domain by recruiting the exocyst, which then captures exocytic vesicles and tethers them to the plasma membrane prior to their fusion (Polgar and Fogelgren, 2018). E-cadherin could also maintain the size of the sub-septate junction domain by organising the sub-membrane F-actin scaffold to anchor membrane proteins and antagonise their endocytosis. This seems less likely, however, since loss of α-catenin, which links E-cadherin adhesion complexes to F-actin, does not lead to a decrease in the size of the domain (Desai et al., 2013).

The expansion of the septate junctions in *shg* and *arm* mutants suggests that E-cadherin adhesion complexes somehow limit their basal extension. This could be by acting as a barrier that prevents septate junction proteins spreading more basally or by promoting their endocytosis at the boundary between the two domains. A third possibility is that septate junction proteins are up-regulated in *shg* and *arm* mutants, leading to lengthening of the septate junctions to compensate for the absence of Cadherin adhesion complexes. As is the case for the sub-septate junction domain, loss of α-catenin has no significant effect on the size of the septate junctions, making it unlikely that the linkage of E-cadherin to F-actin plays a role in establishing the border between the adherens and septate junctions.

N-cadherin does not appear to play any significant role in the formation of junctions in the midgut. N-cadherin loss of function mutants have no defects in apical-basal polarity or junction formation and form septate and sub-septate junction domains of normal size. Furthermore, knock down of N-cadherin does not enhance the phenotype of *shg* mutants, indicating that it does not compensate for E-cadherin in the absence of the latter. In fact, loss of N-cadherin partially suppresses the expansion of the septate junction domain seen upon loss of E-cadherin alone for reasons that are unclear. The only obvious phenotype was an increase in β-catenin levels. Since β-catenin does not detectably localise to the membrane in the sub-septate junction domain in the absence of E-cadherin, N-cadherin does not appear to recruit β-catenin to cell-cell adhesions. It therefore seems likely that N-cadherin somehow destabilises β-catenin, perhaps by activating the APC/axin destruction complex, but this question will require further investigation.

## Methods

### Drosophila stocks

The following *Drosophila melanogaster* stocks were used in this study: *w*^1118^*, FRT42D shg^R69^* (Godt and Tepass, 1998); *FRTG13 shg^IG29^* (Tepass et al., 1996), *hs–Flp FRT40A; da–Gal4 UAS–mCD8::GFP α-Cat1 /TM6B* (Sarpal et al., 2012); *tub–Gal80 ubi–α-Cat FRT40A; act5c–Gal4 α-Cat1 /TM6b*, *FRT19A arm^YD35^* (Riggleman et al., 1989), *FRT19A arm^XP33^* (Riggleman et al., 1989); y w, UAS-mCD8::GFP, Act5C-GAL4, hsFLP[1]; FRTG13 tubP-GAL80 (Lee and Luo, 1999); w, hsFLP, tubP-GAL80, FRT19A;; tubP-GAL4, UAS-mCD8::GFP/TM3, Sb (Lee and Luo, 1999), y w hsFLP tub-GAL4 UAS-GFPnls/FM7; FRT42D tub-GAL80/CyO (Caygill and Brand, 2017), UAS-shg-RNAi (BDSC, 32904), UAS-shg-RNAi (BDSC, 32428), UAS-arm-RNAi (BDSC, 35004), UAS-α-cat-RNAi (BDSC, 33430), UAS-Ncad-RNAi (BDSC, 41982), y w; MyoIA-GAL4, tubP-GAL80[ts] (Chen et al., 2018).

### *Drosophila* maintenance, crosses and genetics

*Drosophila melanogaster* stocks were maintained on standard medium supplemented with dry yeast at 18°C or room temperature (∼22°C) prior to experimental procedures. We used the Mosaic Analysis with Repressible Cell Marker (MARCM) technique to generate mutant clones marked by GFP expression within otherwise heterozygous midguts (Lee and Luo, 1999). Crosses were grown at 25°C for 7 (prepupa stage) or 13 days (mated 3-old day adults) and subsequently subjected to heat-shock. Heat-shock was performed at 37°C for 1 hour, twice per day, for 2 to 5 days. Flies were frequently transferred to new vials to decrease population density and exchange the medium. After heat-shock, flies were transferred to 25 °C until dissection. The female flies used in the experiments were always kept with male siblings and their guts dissected 3 to 16 days after the last heat-shock. When expressing RNAi constructs in clones using the MARCM technique, crosses were maintained at 18°C and placed at 29°C after heat-shock until dissection. For RNAi experiments driven by MyoIA-GAL4, crosses were grown at 18°C, and 3-day old adults were transferred to 29°C for 7-12 days before dissection.

### Immunofluorescence and microscopy

Midgut dissections and immunostainings was performed as described in (Chen et al., 2018) with the following modifications. Female midguts were dissected in a depression slide containing 1XPBS for a maximum of 5 minutes. Dissected midguts were transferred to a 3-10 cm plastic cylinder sealed with a wire mesh and submerged in 1XTSS solution (0.03% Triton X-100, 4 g.L^-1^ NaCl; 95°C) at 78°C for 14 seconds followed by 1 min in 1XTSS solution on ice. Heat-fixed guts were then transferred to an Eppendorf tube containing 8% PFA in 1XPBST (0.1%Triton X-100) and fixed for 20 min at room temperature. Guts were washed 3Xs (10 min each) with 1XPBST and subsequently blocked with 2% Normal goat Serum (NGS) in 1XPBST for 1 hour at room temperature. Midguts were incubated with primary antibodies in 2% NGS in 1XPBST overnight at 4°C. The primary antibodies used in this study were: anti-Arm (DSHB, mouse 1:100), anti-Mesh (gift from M. Furuse, rat 1:1000), anti-Scrib (gift from C. Q. Doe, rb 1:1000), anti-Canoe (gift from M. Peifer, rb 1:1000), anti-Tsp2A (gift from M. Furuse, rb 1:1000), anti-Coracle (DSHB, mouse 1:100), anti-Lgl (SCBT, rb 1:250), anti-aPKC (SCBT, rb 1:100), anti-Nrv (DSHB, mouse 1:100) and anti-GFP (AbCam, chicken, 1:1000). Midguts were washed 3Xs, 20 min each, at room temperature with 1XPBST before incubation with secondary antibodies in 2%NGS in 1XPBST for 2 h at RT. The secondary antibodies used in this study were: 488 anti-mouse (#A11029, 1:1000), 488 anti-rabbit (#A11034, 1:1000), 488 anti-chicken (#A11039, 1:1000), 555 anti-rat (#A21434, 1:1000), 555 anti-mouse (#A21422, 1:1000), 555 anti-rabbit (#A21428, 1:1000), 647 anti-mouse (#A21236, 1:500), 647 anti-rabbit (#A21245, 1:500), and 647 anti-rat (#A21247, 1:500). After incubation with secondary antibodies, the midguts were washed 3Xs, 20 min each, with 1XPBST and afterwards mounted in Vectashield containing DAPI (Vector Laboratories). Images were acquired on an Olympus IX81 confocal microscope using FluoView FV1000 Laser Scanning software (4.2.1.20). Fixed midguts were imaged with a 60X oil objective (UPLSAPO, NA=1.35) in sequential mode (line) with the saturation-warning LUT on. Z-stacks of guts were acquired from the top until the middle section to reveal the enterocytes (ECs) longitudinal profile at 800X800 pixels per image (pixel size=0.265µm) with a 1µm step size between frames with a 12-bit depth. Images were processed using ImageJ (Schindelin et al., 2012).

### Quantification of EC length and circularity

The ImageJ plugin segmented line tool was used to measure the length of the lateral domain. The “total lateral length” corresponds to the distance between the start of the Septate junctions on the apical side until the basement membrane of EC cells (excluding the muscle layer). The “Septate Junction length” corresponds to the total length of the given septate junction marker, while the “Sub-septate junction length” corresponds to the difference between the “Septate junction length” and the “Total lateral length”. Each measurement was carried out in the most longitudinal view of each septate junction marker in cells located on the middle section of the gut. To quantify EC circularity, we used the polygon tool in ImageJ, where the basal area of each EC cell was manually delineated using either the GFP from the MARCM system or lateral membrane marker staining. The value of 1 represents a perfect circle.

### Quantification of fluorescent intensity

To plot Arm intensities across the Adherens junction in Figure 2, a straight line was drawn between two nuclei centered at the Ajs. To quantify Armadillo intensity at the level of the Adherens junctions, we used the segmented line tool with a width of 3. In heterozygous ECs, the AJs were identified by Armadillo staining, whereas in the homozygous ECs, cell-cell borders are identified either by the GFP from the MARCM clones or by the relative position of a septate Junction marker. Line segments were transferred to the Armadillo channel to measure the intensities. To quantify cytoplasmic Armadillo intensity, an oval shape of fixed size – equal for heterozygous and homozygous ECs in each image - was drawn between the lateral domain and the nucleus and measured in the Armadillo channel. Integrated intensity values were used for graphic representation.

### Normalization, statistics and experimental reproducibility

To compare measurements between clones of different mutants, the homozygous mutant individual cell measurements (length of the lateral domains or Armadillo intensities) were first averaged, and the resulting mean divided by the average of the individual measurements in heterozygous enterocytes from the same sample. All controls (heterozygous enterocytes in each clone) were standardized to 1. SuperPlots were used in all graphic visualizations except when directly comparing clones that are represented by a single measurement (Lord et al., 2020). The Shapiro-Wilk test was used to assess normality in all distributions. The Kolmogorov-Smirnov test was used in Figures S1B, S1E, S1F, S3D, 4D, 4E (*shg^R69^* and *arm^YD35^*), 4F (*α-cat^1^*and *arm^YD35^*), S4B (“Septate junction length”), S4D (“Septate junction length”), S4E, 5C, 5D (“Septate junction length”) and 6D; Unpaired t-tests with Welch’s correction were used in Figures S1C, 4E (*α-cat^1^*), 4F (*shg^R69^*), S4B (“Sub-septate junction length” and “Total lateral length”), S4D (“Sub-septate junction length” and “Total lateral length”) and 5D (“Sub-septate junction length” and “Total lateral length”); The Kruskal-Wallis with Dunn’s post hoc test was used in Figures 2G and 2H; A one-way Anova with Tukey’s post hoc test was used in Figure 6C. The mean±SD is represented in black in each distribution and statistical significance is considered at p<0.05. GraphPad prism (10.0.3) was used to build the graphics and to perform statistical analyses. The number of individual experiments/crosses per condition is as follows: UAS::shgRNAi (32428), 1 experiment; UAS::shgRNAi (32904), 2 experiments; UAS::armRNAi(35004), 1 experiment; UAS::α-catRNAi (33430), 2 experiments; UAS::NcadRNAi (41982), 1 experiment; *α-cat^1^*, 4 experiments; *arm^YD35^*, 5 experiments; *arm^XP33^*, 2 experiments; *Ncad^405^*, 2 experiments; *shg^IG29^*, 5 experiments; *shg^R69^*, 5 experiments; *shg^IG29^*, NcadRNAi, 3 experiments.

## Acknowledgements

We would like to thank Mikio Furuse, Chris Doe, Mark Peifer and Ulrich Tepass for fly stocks and antibodies, Jia Chen for help and advice on midgut preparation and staining and the lab members of the Daniel St Jonston lab for technical support. This work was supported by a Wellcome Trust Principal Fellowship to DStJ (080007, 207496) and by core support from the Wellcome Trust (092096, 203144) and Cancer Research UK (A14492, A24823). CAC was supported by a scholarship from the Fundação para a Ciência e Tecnologia, Portugal (PD/BD/52194/2013) and MT by an EU Marie Skłowdowska Action postdoctoral fellowship (GA797837).

## Authors contributions

Conceptualization: CAC, DStJ; Methodology: CAC, MT; Investigation: CAC, MT; Formal analysis: CAC; Resources: DStJ; Data curation: CAC, MT; Visualization: CAC, DStJ; Supervision: DStJ; Funding acquisition: CAC, MT, DStJ; Writing - Original draft preparation: CAC, DStJ; Review and editing: CAC, MT, DStJ.

---

**Figure S1:**
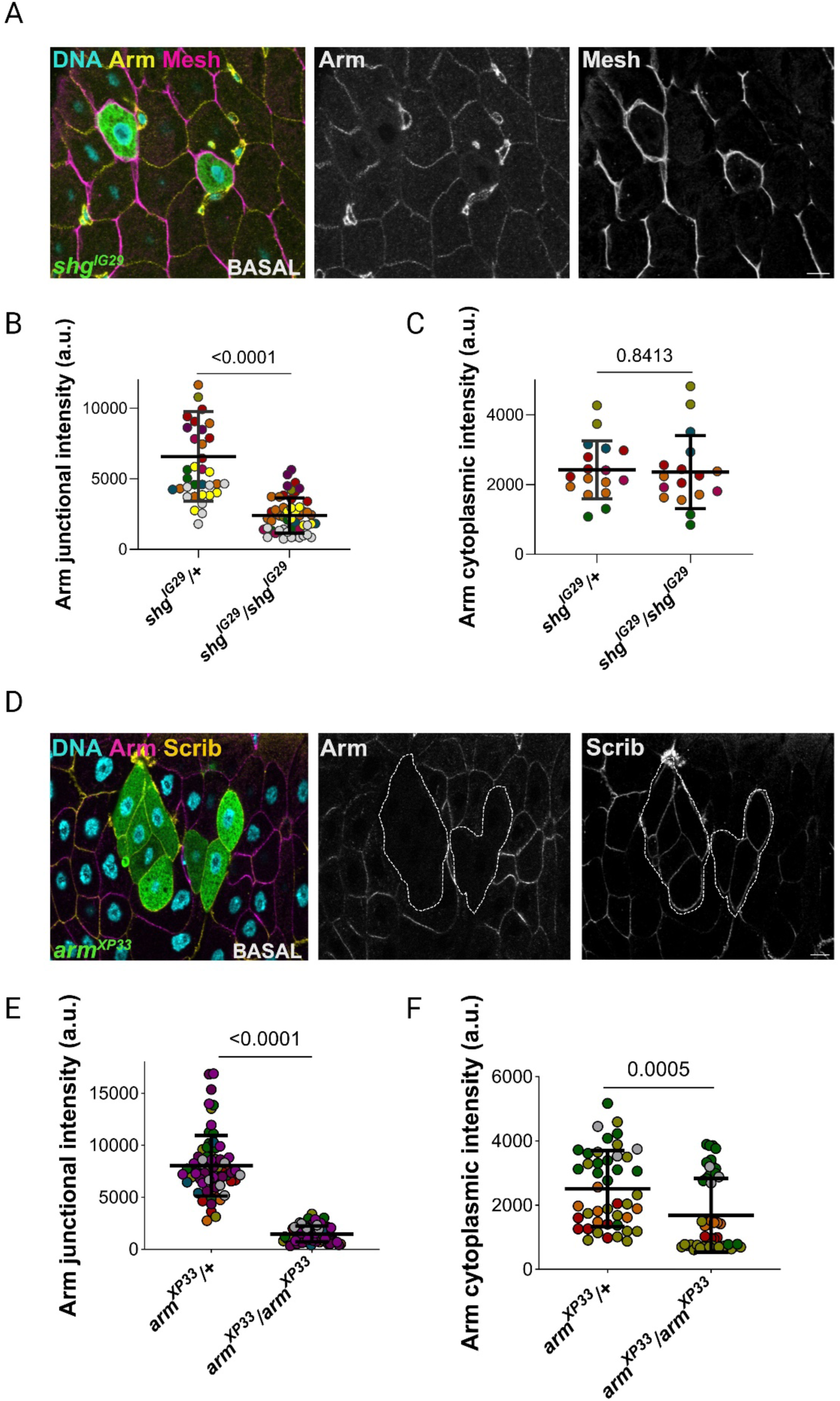
β-catenin is degraded in *shotgun* mutants. **A.** A horizontal confocal section near the basal side of a midgut containing a *shg^IG29^* homozygous clone (GFP; green), stained for DNA (DAPI; blue), Armadillo (magenta) and Mesh (yellow). **B.** Measurements of Armadillo intensity at cell-cell junctions in *shg^IG29^/+* (mean=6577±3183, 38 cell-cell junctions) and *shg^IG29^/shg^IG29^* (mean=2397±1250, 53 cell-cell junctions) enterocytes; **C.** Measurements of Armadillo cytoplasmic intensity in *shg^IG29^/+* (mean=2425±832, 17 cells) and *shg^IG29^/shg^IG29^* (mean=2360±1047, 17 cells) enterocytes; **D.** A horizontal confocal section near the basal side of a midgut containing an *arm^XP33^* homozygous clone (GFP; green), DNA (DAPI; blue), Armadillo (magenta) and Scribble (yellow). **E.** Measurements of Armadillo intensity at cell-cell junctions in *arm^XP33^/+* (mean=8037±2920, 67 cell-cell junctions) and *arm^XP33^/arm^XP33^* (mean=1487±752, 73 cell-cell junctions) enterocytes; **F.** Measurements of Armadillo cytoplasmic intensity in *arm^XP33^/+* (mean=2512±1183, 44 cells) and *arm^XP33^/arm^XP33^*(mean=1684±1144, 49 cells) enterocytes. The scale bars in A and C = 10 µm.

**Figure S2:**
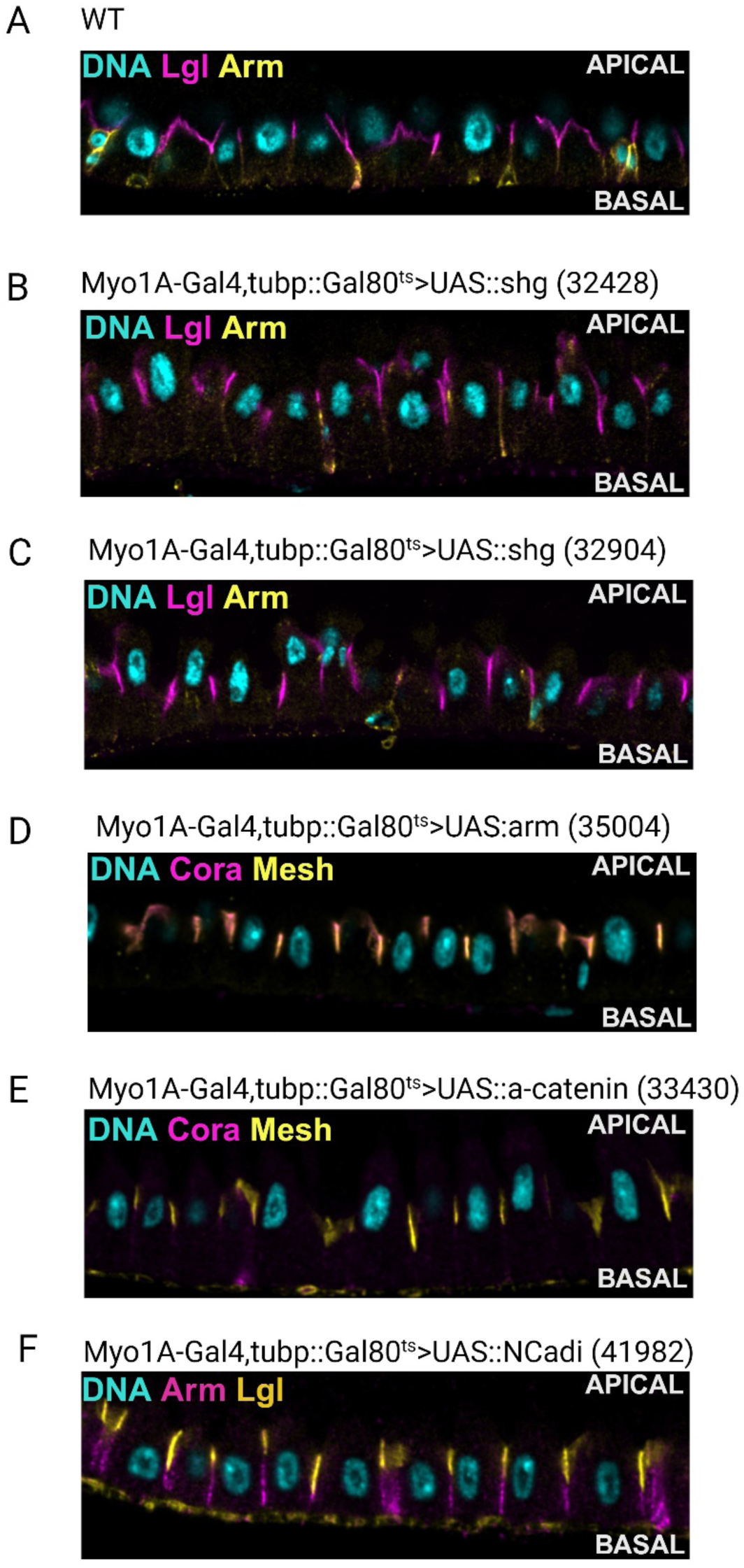
Adherens junctions are not required for the maintenance of apical-basal polarity in the midgut. **A.** An apical-basal section through a wild-type midgut stained for DNA (DAPI; blue), Lgl (magenta) and Armadillo (yellow). **B and C.** Apical-basal sections through midguts expressing *shotgun* RNAi under the control of Myo1A-Gal4, stained for DNA (DAPI; blue), Lgl (magenta) and Armadillo (yellow). **D.** An apical-basal section through a midgut expressing *armadillo* RNAi and stained for DNA (DAPI; blue), Coracle (magenta) and Mesh (yellow). **E.** An apical-basal section through a midgut expressing *α-catenin* RNAi stained for DNA (DAPI; blue), Coracle (magenta) and Mesh (yellow); **F.** An apical-basal section through a midgut expressing *N-cadherin* RNAi stained for DNA (DAPI; blue), Armadillo (magenta) and Lgl (yellow).

**Figure S3:**
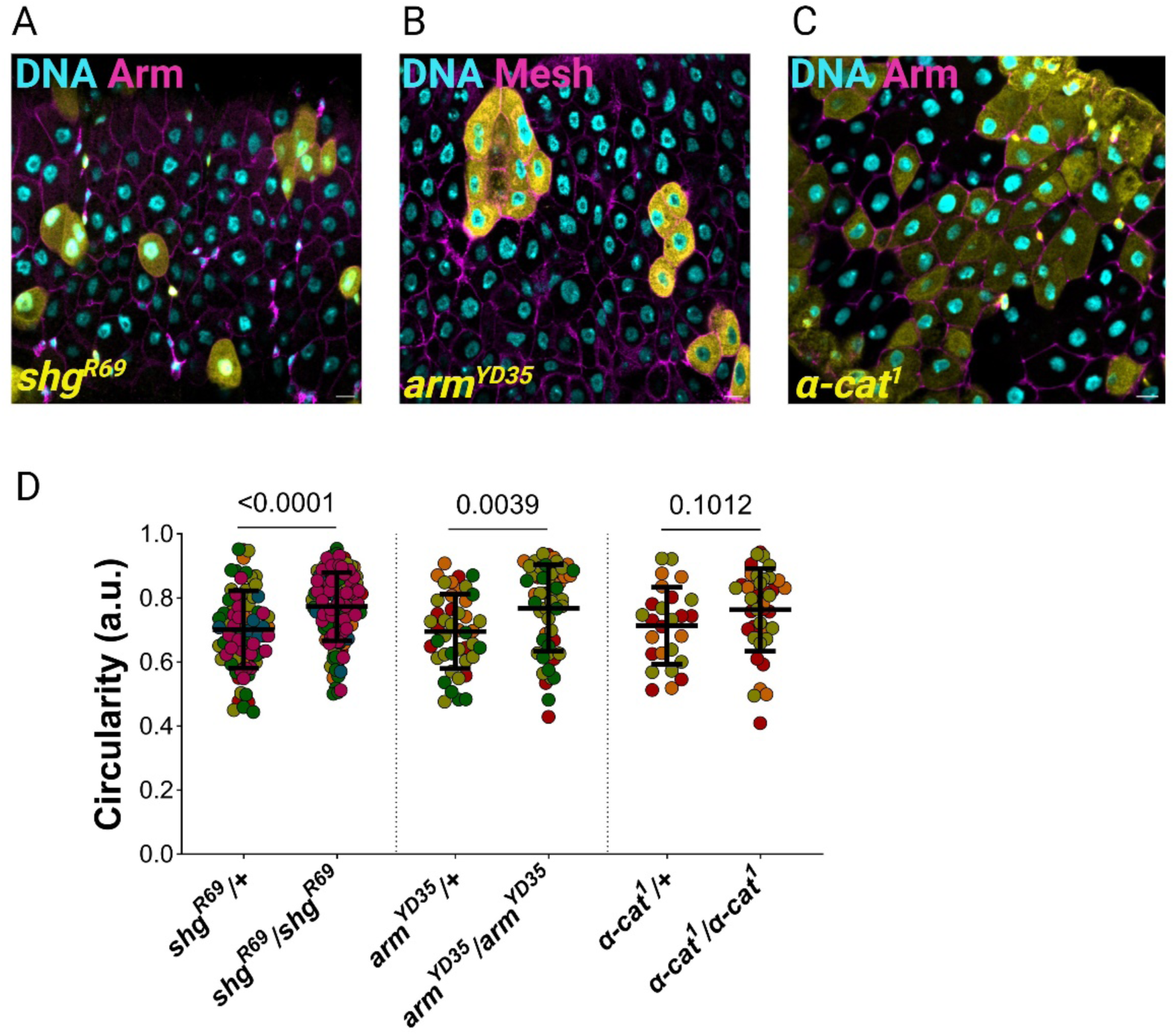
E-cadherin and β-catenin regulate enterocyte shape independently of α-catenin. **A.** Horizontal sections though midguts containing *shg*^R69^ homozygous clones marked by GFP expression (yellow) stained for DNA (DAPI, blue) and Armadillo (magenta). **B.** *arm^YD35^* homozygous clones (yellow) stained for DNA (DAPI, blue) and Mesh (magenta). **C.** *α-catenin*^1^ homozygous clones (yellow) stained for DNA (DAPI, blue) and Armadillo (magenta). Scale bar=10 µm. **D.** Measurements of basal circularity for *shg^R69^/*+ (82 cells), *shg^R69^/shg^R69^* (129 cells), *arm^YD35^/+* (44 cells), *arm^YD35^/arm^YD35^*(52 cells), *α-cat^1^/+* (26 cells) and *α-cat^1^/α-cat^1^* (40 cells) enterocytes. The scale bars = 10 µm.

**Figure S4:**
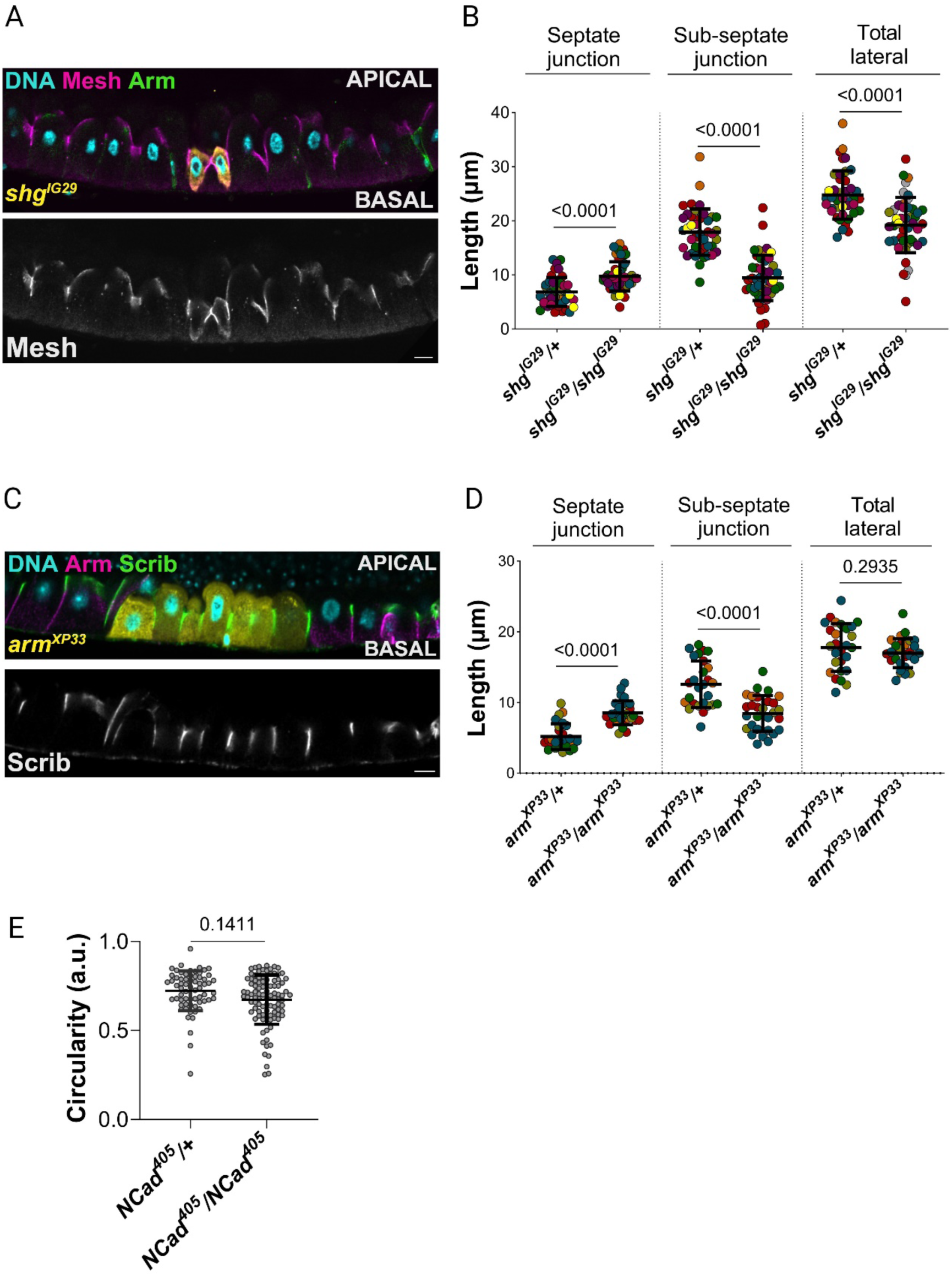
E-cadherin and β-catenin regulate Septate junction length. **A.** An apical-basal section of a midgut containing homozygous cells mutant for a second *shotgun* allele, *shg^IG29^* (yellow) stained for DNA (DAPI, blue) and Mesh (magenta). **B.** Measurements of the lengths of the septate junctions (*shg^IG29^/+*: 6.86±2.67µm, 43 junctions; *shg^IG29^/shg^IG29^*: 9.75±2.70µm, 46 junctions), the sub-septate junction domain (*shg^IG29^/+*: 17.91±4.29µm, 43 junctions; *shg^IG29^/shg^IG29^*: 9.47±4.20µm, 46 junctions) and the total lateral length (*shg^IG29^/+*: 24.77±4.47µm, 43 junctions; *shg^IG29^/shg^IG29^*: 19.22±5.11µm, 46 junctions) in *shg^IG29^* heterozygous and homozygous enterocytes; **C.** An apical-basal section of a midgut containing a homozygous mutant clone for a second *arm* allele *arm^XP33^*(yellow) stained for DNA (DAPI; blue), Armadillo (magenta) and Scribble (yellow). **D.** Measurements of the lengths of the septate junctions (*arm^XP33^/+*: 5.19±1.82µm, 27 junctions; *arm^XP33^/arm^XP33^*: 8.54±1.70µm, 29 junctions), sub-septate junction domain (*arm^XP33^/+*: 12.59±3.30µm, 27 junctions; *arm^XP33^/arm^XP33^*: 8.45±2.53µm, 29 junctions) and the total lateral length (*arm^XP33^/+*: 17.78±3.37µm, 27 junctions; *arm^XP33^/arm^XP33^*: 16.98±2.03µm, 29 junctions) in *arm^XP33^/+* and ; *arm^XP33^/arm^XP33^*enterocytes. **E.** Measurements of the circularity of *Ncad^405^/*+ (65 cells) and *Ncad^405^/ Ncad^405^* (96 cells) enterocytes. Scale bars = 10 µm.

